# Nanoparticle-based local translation reveals mRNA as translation-coupled scaffold with anchoring function

**DOI:** 10.1101/483727

**Authors:** Shunnichi Kashida, Dan Ohtan Wang, Hirohide Saito, Zoher Gueroui

**Affiliations:** PASTEUR, Département de chimie, École normale supérieure, PSL University, Sorbonne Université, CNRS, 75005 Paris, France; Institute for Integrated Cell-Material Sciences (iCeMS), Kyoto University, Yoshida-Honmachi, Sakyo-ku, Kyoto, 606-8501, Japan; The Keihanshin Consortium for Fostering the Next Generation of Global Leaders in Research (K-CONNEX), Yoshida-Honmachi, Sakyo-ku, Kyoto-shi, Kyoto, 606-8501, Japan; Department of Life Science Frontiers, Center for iPS Cell Research and Application, Kyoto University, 53 Kawahara-cho, Shogoin, Sakyo-ku, Kyoto, 606-8507, Japan

## Abstract

Spatial regulations of mRNA translation are central to cellular functions and relies on numerous complex processes. Biomimetic approaches could bypass the endogenous complex processes, improve our comprehension, and allow for controlling local translation regulations and functions. However, the causality between localizing translation and nascent protein function remains elusive. Here, we develop a novel nanoparticle-based strategy to magnetically control mRNA spatial patterns in mammalian cell extracts and investigate how local translation impacts nascent protein localization and function. By monitoring translation on magnetically localized mRNAs, we show that mRNA-nanoparticle operates as a source for the continuous production of proteins from defined positions. By applying magnetic localization of mRNAs coding for Actin Binding Proteins, we trigger the local formation of actin cytoskeleton and identify minimal requirements for spatial control of actin filament network. In addition, our bottom-up approach identifies a novel role of mRNA as translation-coupled scaffold for nascent N-terminal protein domain functions. Our approach will serve as a novel platform for regulating mRNA localization and investigating a functional role of nascent protein domains during translation.

## Introduction

One essential element often missing to explain cell-fate control concerns the spatiotemporal regulation of gene expression. Though DNA stand compacted in nucleus, mRNA species are exported from nucleus and transported to precise locations in cytosol before being translated. Translational control together with the localization of mRNAs significantly contributes to many important cellular processes, such as the establishment of polarity, asymmetric division, synaptic plasticity, and memory consolidation(*1*). For instance, the local translation of actin-associated proteins from subcellular localized mRNAs has been increasingly recognized as an important process for regulating the dynamic formation of actin cytoskeleton during cell polarization(*2*–*7*). Although biochemistry, genetics, cell imaging, biophysics, and “omics” approaches have increased our understanding of the molecules involved and reaction mechanisms of mRNA localization and local translation, none of them can distinguish the functional contribution by spatiotemporal dynamic control from that by molecule-based signalling. Thus, the causality between localizing translation and cell function remains elusive. Being able to design a bottom-up approach to bypass the complex subcellular transport systems and uncouple endogenous biochemical regulation from physical constraints could reveal the principles underlying complex biological systems. *In vitro* synthetic biology and cell-free systems have been powerful approaches for understanding the physical constraints that can shape gene expression networks such as molecular crowding, compartmentalization, and space (diffusion, gradient)(*8*–*12*).

Nevertheless, having access to the spatiotemporal nature of translation is still challenging and requires the development of novel orthogonal methods for harnessing biomolecule functions. Optogenetics, magnetogenetics, and nanoparticle-based technologies could provide spatiotemporal control of biomolecule activities, but these approaches remain mainly limited to control signalling pathways and transcription-based gene expression(*13, 14, 23*–*32, 15*–*22*) and have not yet achieved spatiotemporal translation control.

Here, we examined how magnetic control can be implemented to spatially manipulate mRNA localization and its translation without introducing genetic interference to the cell. As first step towards this goal, we developed an *in vitro* local translation model based on synthetic-mRNA conjugated magnetic nanoparticles to control translation-based protein synthesis in space within confined *HeLa* cell extracts (**Figure 1**). The mRNA-nanoparticle complexes (mRNA-NPs) were designed as movable mRNA-based protein factories upon magnetic control. When supplied with sufficient translational ingredients, they enable continuous syntheses of specific proteins with spatiotemporal control (**Figure 1**). We achieved spatial patterns of mRNA-NPs in confined *HeLa* cell extracts using magnetic forces to trigger asymmetric mRNA-NP string-like patterns (**Figure 1**). Using this platform, we simultaneously monitored localized translation and the dynamic behaviour of the protein products. The results revealed an unexpected role of mRNA as an anchoring molecule to create a local concentration and function of nascent protein domains during translation. Its functional relevance to cell function is demonstrated by the striking differences in actin filament patterns that were triggered by simply rearranging an Actin Binding Protein domain toward the N-terminal or the C-terminal end of the protein translated on the mRNA-NPs.

**Figure 1.**
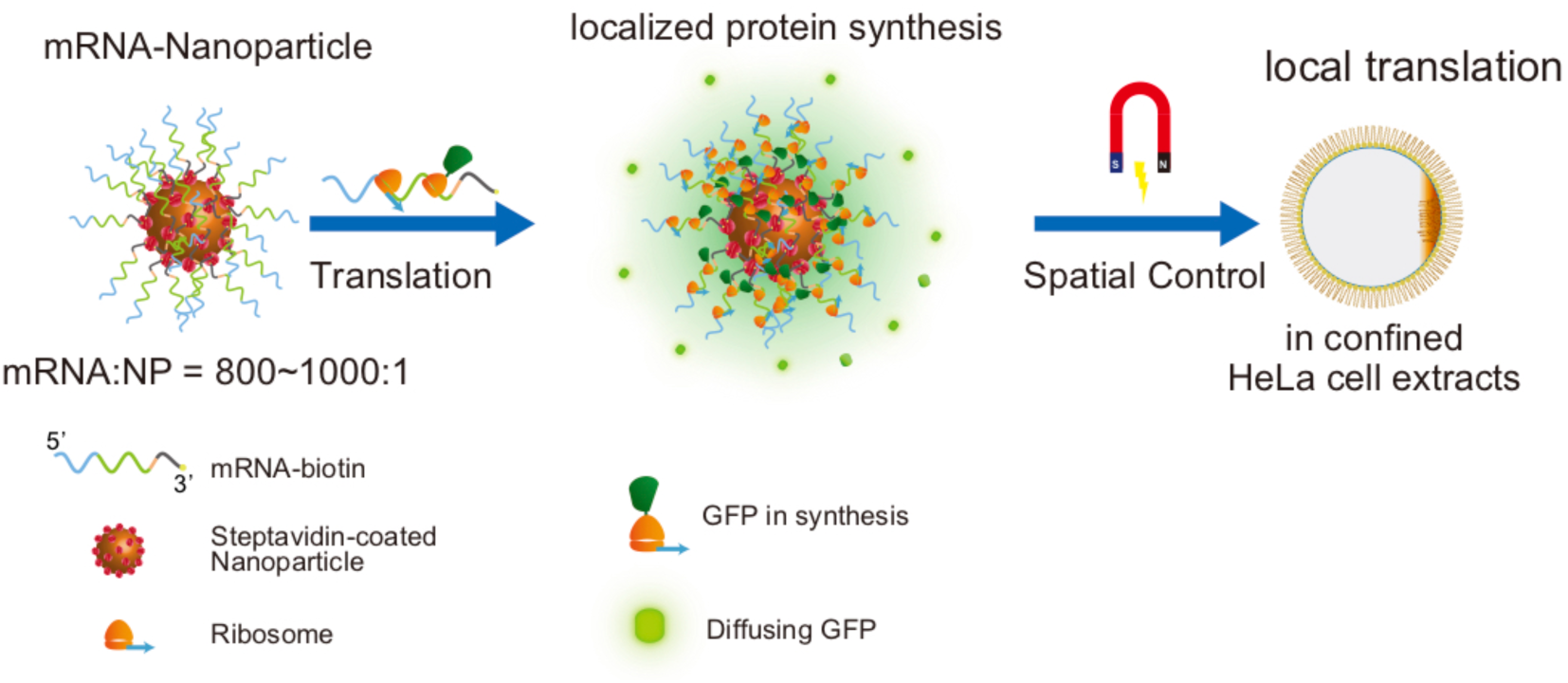
Magnetic nanoparticle-based spatiotemporal regulation of mRNA translation. Messenger RNAs conjugated to iron-oxide nanoparticles (mRNA-NPs) are designed as RNA-based local source of protein synthesis which combine features of magnetic mobility and specific mRNA localization around nanoparticle. The mRNA-NPs can continuously synthesize specific nascent proteins (e.g., GFP) and release them from the nanoparticles. Spatial Control of mRNA-NPs is mediated by magnetic field to generate mRNA localization in order to assess their effect on local translation.

## Results

### Translation of mRNA - nanoparticle conjugates in *HeLa* cell extracts

As a first step towards the spatial manipulation of mRNA molecules using magnetic forces, we designed mRNA conjugated magnetic nanoparticles (mRNA-NPs) and assessed their translation efficiency by monitoring GFP protein synthesis in *HeLa* cell extracts. Messenger RNAs encoding green fluorescent protein (GFP) were transcribed *in vitro* and biotinylated at their 3-prime end. The mRNAs were grafted onto the surface of streptavidin conjugated superparamagnetic nanoparticles (size 200 nm, see methods) by mixing in high salt buffer at a stoichiometry of about 1650 mRNAs per NP (**Figure S1a**).

We next monitored GFP protein synthesis from mRNA-NPs in *HeLa* cell extracts, a model cytoplasm with a full complement of translational factors. To mimic the geometrical confinement by cell membranes, the extracts and mRNA-NPs were encapsulated in spherical droplets dispersed in mineral oil with diameters ranged between 10-130 micrometres (methods, **Figure S1b**). To assess the translational efficiency of the mRNA-NPs, we monitored the fluorescence of GFP translated from GFP-encoding mRNA conjugated with NPs in encapsulated extracts and found that GFP proteins were continuously produced from mRNA grafted on nanoparticles as function of time (**Figure S2**). Thus, mRNA grafted on nanoparticles does not affect mRNA translatability and mRNA-NPs can be used for studying translation and protein synthesis.

Next, we assessed whether we could observe the asymmetrically localized mRNA-NPs and its translation efficiency in the droplets. For fluorescent detection of mRNA-NPs, Cy5-labeled biotin-mRNAs were supplemented with non-fluorescent biotin-mRNA during the conjugation with NPs (Methods). Droplets of cell extracts were transferred into a slide chamber at the vicinity of a permanent NdFeB magnet that generates a field of about 0.15 T and a gradient of 50 T.m^-1^ (Methods, **Figure 2a**). Within a few seconds after applying magnetic field, the dispersed mRNA-NPs became organized into micrometric string-like assemblies that were aligned perpendicularly to the magnet side of the droplet interface (**Figure 2b**). The droplets were incubated for 4 hours with or without magnet condition and monitoring their fluorescent intensity showed that translated GFP were homogeneously distributed within the droplets **(Figure 2c and S2b)**. This suggests that the newly synthesized GFP proteins were successfully released from mRNA-NPs and rapidly diffused throughout encapsulated cell extracts. Relative GFP concentration showed that the efficiency of GFP synthesis was comparable with or without magnetic field, indicating that neither the magnetic field nor the string-like mRNA-NP assemblies perturbed the translation process **(Figure 2c and S2b)**.

**Figure 2.**
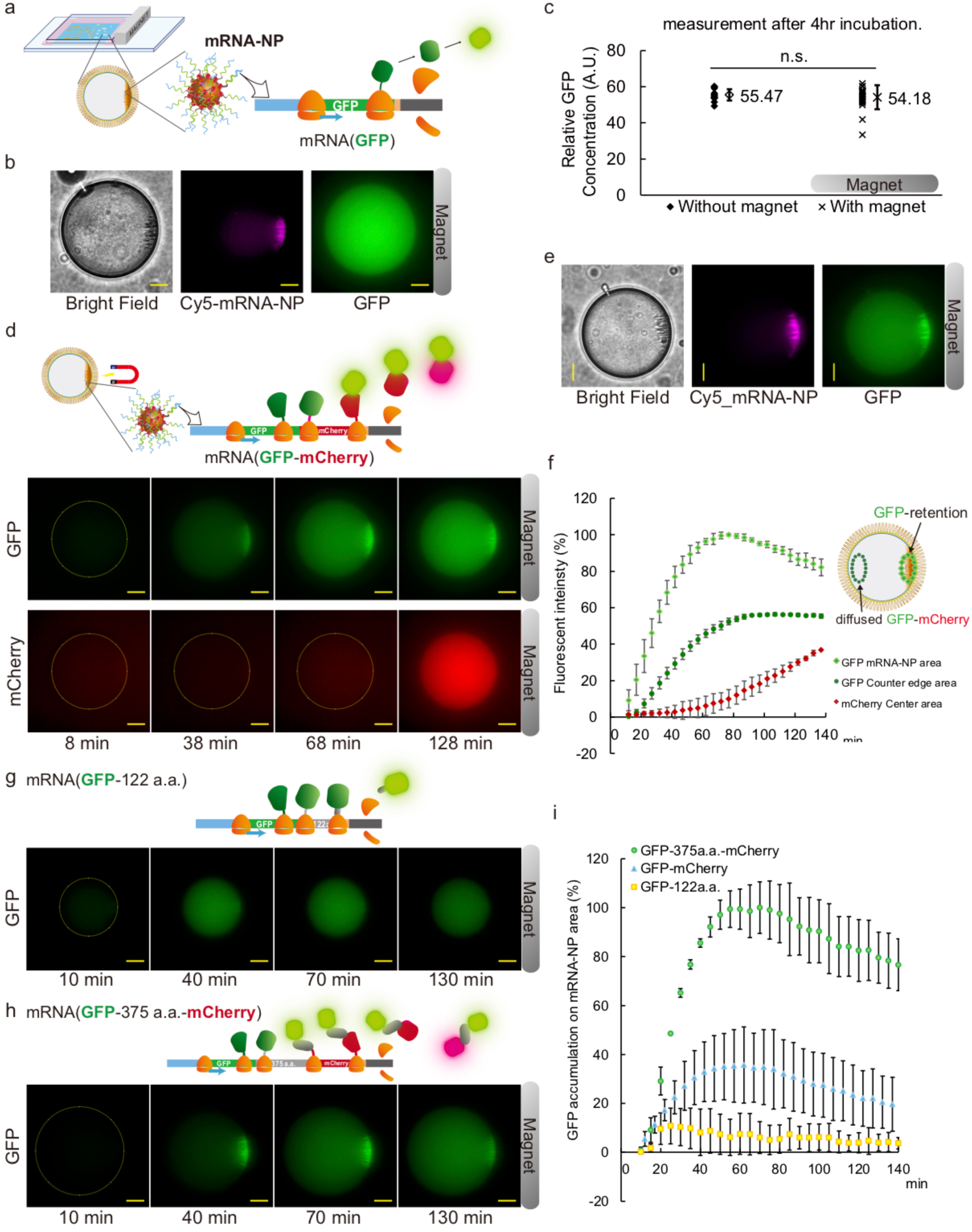
Translation of mRNA - nanoparticle conjugates in HeLa cell extracts droplets. **a-c** GFP translation from localized mRNA-NPs. **a**, Schematic of the translation from asymmetrically localized mRNA-NPs performed in *HeLa* cell extract confined in droplets. The *HeLa* extracts droplets were incubated in an observation chamber beside a permanent magnet. **b**, Epifluorescence microscopy observation of Cy5 labeled mRNAs conjugated to NPs and asymmetrically aligned perpendicularly to the magnet side of the droplet interface. Translation of mRNA-NPs produces GFP that are homogeneously diffusing within the droplet. Scale bar, 20 µm (**b**). **c**, Relative GFP concentration measured in droplets after 4hr incubation with or without magnet. **d-f** GFP-mCherry translation from asymmetrically localized mRNA-NPs. **d**, Top: Schematic of translation process on localized mRNA-NPs: N-terminal nascent GFP are anchored to mRNA-NP via translating mCherry peptides and ribosome. Bottom: Representative time-lapse images of translated GFP and mCherry with epifluorescence microscopy. **e**, Cy5 labeled mRNA on NPs are colocalizing with translated GFP. **f**, Time-dependent evolution of the normalized mean fluorescent intensity of GFP and mCherry in selected regions of interested of the droplets (one frame every 5 minutes). Error bars represent standard deviation (SD). **g,h,** Representative time-lapse images of the translated GFP from localized mRNA-NPs for mRNA(GFP-122a.a) and mRNA(GFP-375a.a.-mCherry), respectively. **i,** Time dependent evolution of the mean fluorescent intensity of accumulated GFP on mRNA-NPs for mRNA(GFP-375a.a.-mCherry), mRNA(GFP-122a.a) and mRNA(GFP-mCherry).

### Monitoring of nascent proteins on localized mRNA-NPs

To better characterize mRNA-NP complexes, we developed an assay to monitor protein synthesis on localized mRNA-NPs and designed a fusion protein permitting the use of the nascent N-terminal GFP as real-time reporter of local translation. We hypothesized that the translation duration of GFP-mCherry would be extended to about twice compared with that of GFP and will allow nascent GFP to monitor the location of its own mRNA until release of GFP-mCherry proteins (**Figure 2d scheme)**. To investigate this hypothesis, the translation of mRNA(GFP-mCherry)- NPs was analysed using the time-lapse microscopy. About 10 minutes after reaction started, we first observed GFP signal accumulation that was perfectly matching the localized mRNA-NP pattern (**Figure 2e**). Alongside, the GFP fluorescence within the droplets increased homogeneously, indicating that GFP-mCherry proteins were continuously translated, and released from the mRNA-NPs (**Figure 2d, 2f, S3 and Supplementary Movie 1 & 2**). After 70-80 minutes, the GFP signals on the mRNA-NPs reached a maximum and then slowly decayed, possibly because of the limitation of translational efficiency of confined cell extracts. Regarding mCherry, fluorescence started to increase homogeneously within the droplets 50 minutes after reaction was initiated, in agreement with the slower mCherry maturation time (40 min) compared to GFP(*33*–*35*) (8 min, **Figure 2f, red curve and Supplementary Movies 1 & 2**). In addition, we did not observe any co-localization between mCherry and mRNA-NPs, confirming that GFP-mCherry fusion proteins did not stall or accumulated on NP surfaces.

These observations identified several remarkable properties of local translation mediated by mRNA-NPs. The mRNA-NPs function as local continuous production of GFP-mCherry proteins that are released and dispersed in solution. Then, as we hypothesized, the accumulation of GFP fluorescence on mRNA(GFP-mCherry)-NPs indicates that nascent N-terminal GFP was spatially retained on the mRNA via ribosomes during mCherry synthesis (**see Schematic Figure 2d**). Thus, we could consider the N-terminal GFP as a transient fluorescent indicator of local translation from mRNA-NPs. To further investigate the link between protein synthesis and the spatial retention of nascent protein domain, we next monitored GFP local synthesis as function of the length of the downstream sequence after the GFP sequence. We designed two RNA constructs whose downstream sequences were shorter and longer than the mRNA(GFP-mCherry) respectively: the mRNA(GFP-122aa) and the mRNA(GFP-375aa-mCherry) (**Figures 2g and 2h**). We found a small and short-lived GFP accumulation on RNA-NPs during mRNA(GFP-122aa) translation (**Figures 2g, S3** and **Supplementary Movie 3)**, while GFP protein strongly accumulated on nanoparticles during translation of RNA(GFP-375aa-mCherry)-NPs (**Figures 2h, S3** and **Supplementary Movie 4**). These data underlie that the spatial retention time of GFP on mRNA-NPs is correlated with the peptide elongation length during translation, as showed by the time delay difference between GFP signal on RNA-NPs and the GFP diffuse fraction in the droplet (**Figures 2i** and **S4**). From a biological perspective, this phenomenon may represent a simple translation-based retaining mechanism of proteins at the site of translation for a time span defined by mRNA translational duration time.

### Spatiotemporal control of F-actin meshwork by localized mRNA translation

Next, we asked whether the spatial retention of N-terminal protein domain on localized mRNA-NPs could produce downstream functional diversity. We chose actin cytoskeleton pattern formation because its relevance to important functions of a living cell, its versatile patterns, and its high dynamic nature(*36, 37*). Many efforts have been made on deciphering the molecular players involved in targeting and locally regulating actin-related mRNAs, yet current models are tightly dependent on the cellular contexts through multiple processes and regulators. We focused on a function domain of Actin Binding Proteins widely shared by actin binding protein superfamily. We replaced the mCherry protein of the GFP-mCherry fusion with a functional domain of the Actin Binding Domain of Utrophin (here after called ABD) as functional domain (**Figure 3**), which is shared among a group of the Actin Binding Protein superfamily such as Dystrophin and α-Actinin(*38*–*41*). This 30 kDa ABD contains two actin-binding sites to bind and stabilize actin filaments and its GFP fusion protein can be used as actin filament promoter and label(*42*).

**Figure 3.**
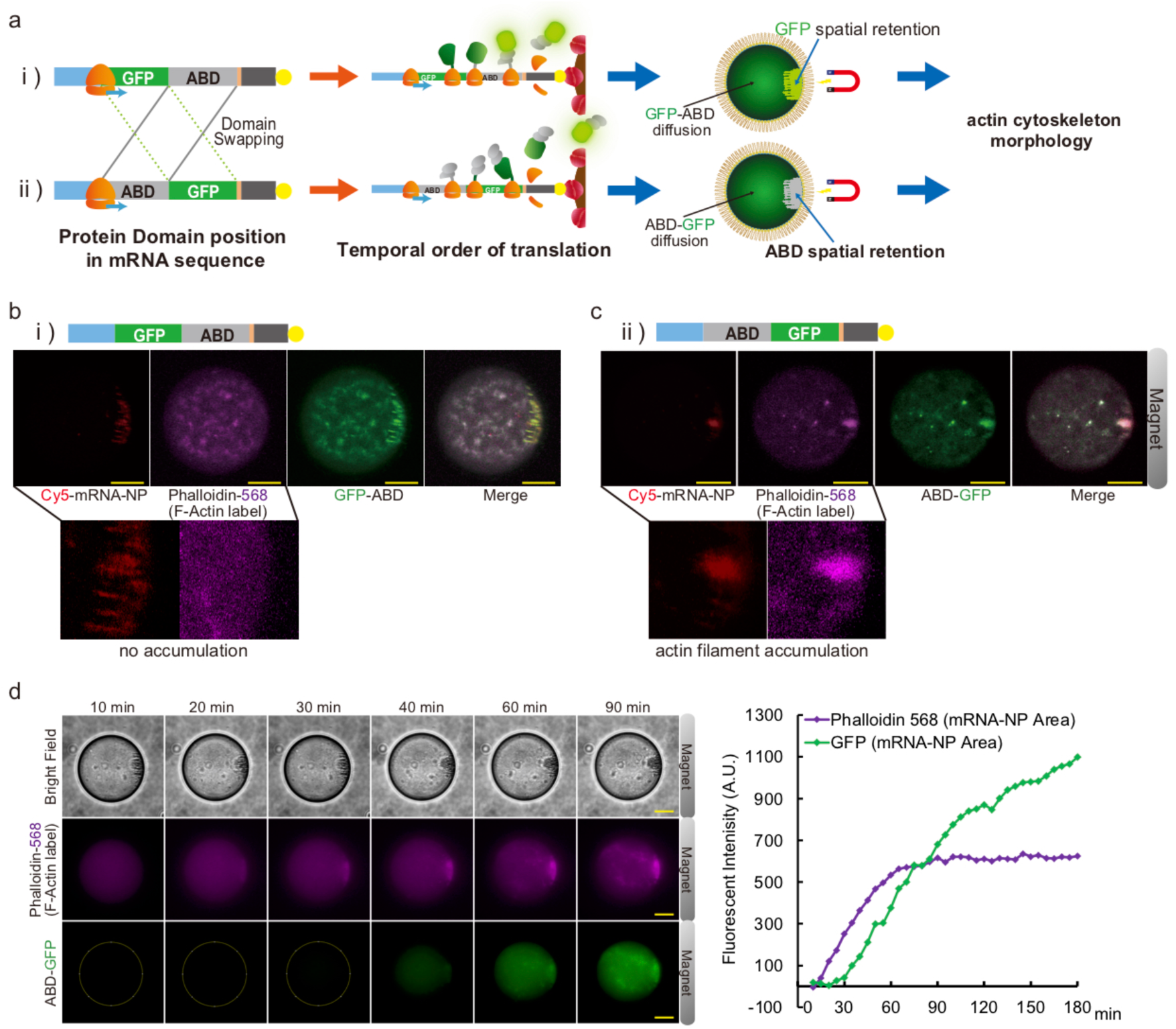
Spatial control of F-actin meshwork by localized mRNA translation highlight the key role of the spatial retention of Actin Binding Protein. **a,** To assessing the effect of the spatial retention of N-terminal Actin Binding Protein (ABD) during local translation on controlling actin filament assembly, two mRNA sequences were designed by swapping the order of ABD and GFP: ABD-GFP and GFP-ABD. Then mRNAs were translated from localized mRNA-NPs in *HeLa* extract droplets. The translation of mRNA(GFP-ABD)-NPs were designed to spatially retain N-terminal GFP on the localized mRNA-NPs, whereas the translation of mRNA(ABD-GFP)-NPs will spatially retain N-terminal ABD on the localized mRNA-NPs. **b, c,** Representative confocal fluorescence microscopy images of Cy5-mRNA-NPs, Phalloidin-568 (F-actin marker), and GFP-ABD (**b**) or ABD-GFP (**c**), after 3 hours of translation. **b, zoom:** formation of a homogenous F-actin meshwork without any local enrichment of filaments at the vicinity of the mRNA-NPs. **c, zoom:** formation of a local and dense F-actin meshwork colocalizing with mRNA-NPs. **d.** Left panel: Representative time-lapse images of F-actin formation upon translation of localized mRNA(ABD-GFP)-NPs. Phalloidin-568 (F-actin marker), and ABD-GFP. Right panel: Quantification of the mean fluorescent intensity of Phalloidin-568 and GFP signal as a function of time in mRNA-NP area. Scale bar, 20 µm (**b-d**).

To investigate the significance of N-terminal position of ABD in mRNA sequence, two mRNA fusions were developed with swapping the order of ABD and GFP (ABD-GFP and GFP-ABD) (**Figure 3a, left**). The mechanism of spatial retention permitted to anticipate two situations: The translation of mRNA(GFP-ABD)-NPs were designed to spatially retain N-terminal GFP on the mRNA-NPs area and will produce GFP-ABD that will diffuse immediately after synthesis, whereas the translation of mRNA(ABD-GFP)-NPs will spatially retain N-terminal ABD on the mRNA-NP area and then release ABD-GFP diffusing homogeneously in the droplets (**Figure 3a**).

First, we checked the translation of these two mRNAs and their capacity to trigger the formation of actin filaments (F-actin). The two mRNAs (without been conjugated to nanoparticles) were translated for 3 hours in cell extract droplets supplemented with monomeric actin and fluorescent phalloidin, a probe for observing F-actin (**Figure S5**). We found that both mRNA constructs generate the formation of an interconnected F-actin meshwork (**Figure S5**). Phalloidin staining was perfectly matching the translated ABD-GFP and GFP-ABD localization, which indicated that their ABDs have the capacity to both promote and bind the F-actin in cell extracts (**Figure S5**).

To assess actin filament formation from localized mRNAs, we next conjugated each mRNA construct with NPs (with 800:1 ratio) and assembled the mRNA-NPs into string-like patterns in confined extracts. Interestingly, when examining the translation of asymmetrically distributed mRNA-NPs, we found that the pattern of F-actin meshworks was highly dependent on the N-terminal or C-terminal position of the ABD sequence on mRNAs (**Figures 3b and 3c**).

First, the translation of localized mRNA(GFP-ABD)-NPs led to the formation of a homogenous F-actin meshwork without any local enrichment of filaments at the vicinity of the mRNA-NPs (**Figure 3b zoom**), as indicated by the phalloidin signal (**Figure 3b**, **Cy5-mRNA-NPs and Phalloidin**). In addition, we also observed the accumulation of GFP signal colocalizing with Cy5-labelled mRNA-NPs and that confirmed the spatial retention of N-terminal GFP during downstream ABD local translation as in the case of GFP-mCherry (**Cy5-mRNA-NP and GFP-ABD in Figures 3b and S6**). In contrast, the local translation from mRNA(ABD-GFP)-NPs showed dense accumulation of Phalloidin signal around the Cy5 labelled mRNA-NPs which suggested that the F-actin formation were locally nucleating and promoting by the spatially retained N-terminal ABD on mRNA-NPs (**Cy5-mRNA-NP and Phalloidin in Figure 3c zoom**). Moreover, string-like assemblies of mRNA-NPs displayed a more compact and denser pattern as if the formation of a F-actin meshwork had modified the mRNA(ABD-GFP)-NP spatial organization (**Cy5-mRNA-NP and Phalloidin in Figures 3b and 3c zoom**). Though some diffused ABD-GFP molecules randomly promoted the F-actin formation in the rest of droplets, the locally translated ABD-GFP mainly co-recruited around mRNA-NPs with accumulated F-actin cytoskeleton (**Cy5-mRNA-NP, Phalloidin, and GFP-ABD in Figure 3c and S6**). The formation of localized F-actin mediated by spatially retained N-terminal ABD was also confirmed using time-lapse microscopy (**Figure 3d and Supplementary Movie 5**). F-actin accumulation on mRNA-NPs formed prior to non-localized actin formation, as revealed by the local enrichment in Phalloidin staining, appearing at early stage of translation (after about 10 minutes, **Figure 3d and Supplementary Movie 5**). The observation of ABD-GFP staining F-actin meshwork was detected latter (after about 40 min), consistently with the timescale of steady-state production of full-length ABD-GFP proteins (**Figure 3d**). Time-lapse observation also showed that the compaction of string-like assemblies of mRNA-NPs into a more compact pattern was correlated with the formation of localized F-actin meshworks (**Figure 3d and Supplementary Movie 5)**.

Altogether these data showed that the N-terminal ABD position in mRNA sequence and mRNA localization in droplet collaboratively dictated the micrometric local formation of F-actin cytoskeleton.

### Spatial Retention of Actin Binding Proteins on hotspot of translational activity promotes the reorganization of actin filament meshwork

Our observations strongly suggest that mRNA(ABD-GFP)-NPs behave as hotspots of active ABD-covered nanoparticles that promote local F-actin formation. In addition, we found that mRNA(ABD-GFP)-NP string-like patterns were changed upon local F-actin formation and eventually formed dense organization. Accordingly, we next examined how the spatial organization of mRNA-NP complexes can be regulated by their own protein products, thus in the absence of external magnetic forces. We first examined how individual mRNA(GFP-ABD)-NPs and mRNA(ABD-GFP)-NPs could trigger different F-actin morphologies. As expected, the translation of mRNA(GFP-ABD)-NPs led to the formation of a homogenous F-actin meshwork with a homogeneously spatial distribution of NPs within the droplets (**Figures 4a** and **4c**), as the one previously observed in **Figure 3b**. Time-dependent confocal observations permit to distinguish the local translation activity on mRNA(GFP-ABD)-NPs, as indicated by the co-localization of Cy5 labelled-mRNAs with GFP fluorescence (**Figure 4a, 4c, S7, and Supplementary Movie 6 top**). In contrast, the translation of mRNA(ABD-GFP)-NPs led to a major spatial reorganization of mRNA-NPs as function of time (**Figures 4b, 4d, S7, and Supplementary Movie 6 bottom**). After 15 minutes, the initially homogeneously distributed mRNA-NPs in the droplets started to form random micrometric clusters and eventually evolved into large assemblies within 45 minutes (**Figure 4d**). Interestingly, this spatial reorganization of mRNA-NPs was correlated with the spatial remodelling of the F-actin meshwork (**Figure 4d**). The formation of a hybrid meshwork composed of F-actin connected to NPs may explain the dynamic clustering of mRNA-NPs observed in Figure 4d and may account for the compaction of mRNA-NP string-like structures into dense organizations (**Figures 3c** and **3d)**. To test this hypothesis, we monitored the translation of non-biotinylated mRNA(ABD-GFP) in the presence of unconjugated NPs (**Figure S8 and Supplementary Movie 7**). In this case, F-actin formed a homogeneous meshwork coexisting with randomly distributed NPs, thus phenocopying the translation of mRNA(GFP-ABD)-NPs (**Figure S8 and Supplementary Movie 7**). Thus, the multivalency provided by the multiple mRNAs grafted to NP and functioning as local hotspots of translational activity is required for the large-scale reorganization of F-actin meshworks.

**Figure 4.**
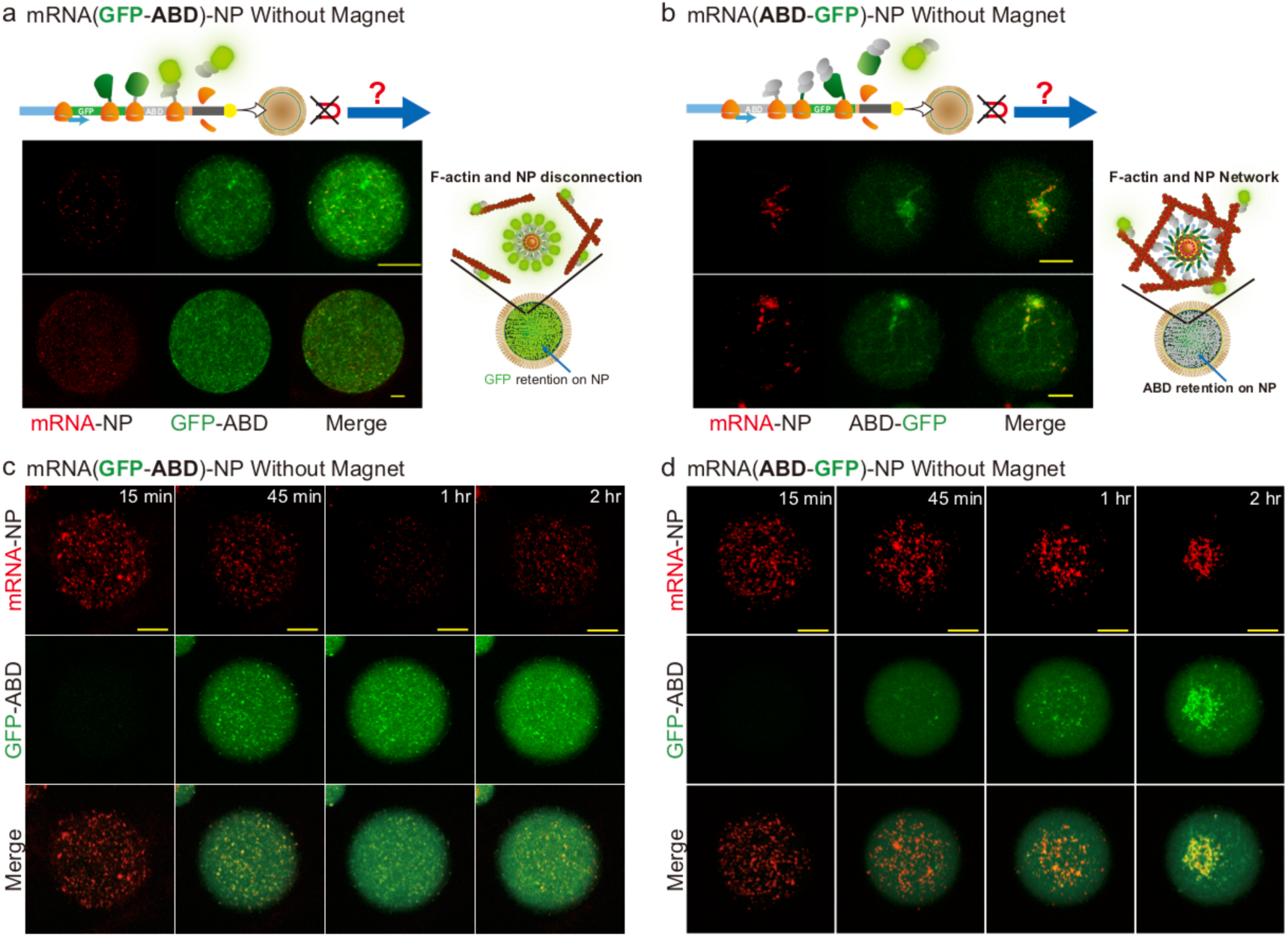
Spatial Retention of Actin Binding Proteins on hotspot of translational activity promotes the reorganization of actin filament meshwork a,b, Confocal observations of translation from mRNA(GFP-ABD)-NPs (**a**) and mRNA(ABD-GFP)-NPs (**b**) to assess how the spatial organization of mRNA-NP complexes can be regulated by their own protein products. Observations were performed after 2-hour translation. F-actin meshwork is labelled by GFP-ABD and mRNA-NPs by Cy5 labelled-mRNAs. **c,d,** Time-lapse observations of the dynamic of mRNA translation and mRNA-NP complexes **c,** N-terminal GFP signals (green) are localized on mRNA(GFP-ABD)-NPs (Red, Cy5 labelled-mRNAs) after 45 minutes of translation. mRNA-NPs are spatially distributed in the droplet and GFP-ABD bound actin filaments form a homogenous meshwork. **d,** Translation of mRNA(ABD-GFP)-NPs lead to a spatial reorganization of mRNA-NPs as function of time: mRNA(ABD-GFP)-NPs (Red) and ABD-GFP (green) bound to actin filaments spontaneously accumulate together to form a dynamic meshwork that is remodeling spatially. Scale bar, 20 µm.

Altogether these data suggest that the spatial retention of ABD on hotspot of translational activity impact the spatiotemporal dynamics of F-actin assemblies, which can, in turn, regulate the localization of the mRNA-NP complexes. Thus, the temporary retention of ABD renders the mRNA complexes interactive with its own coding protein and downstream functional products.

## Discussion

Our bottom-up approach is a first step towards the spatial control of mRNA translation using magnetic nanoparticles, thus expanding the applications of magnetogenetics and nanoparticle-based technologies to RNA-based gene expression and spatial control. The designed mRNA-based complexes provide a novel platform allowing the spatial manipulation of local translation. These functionalities relied on two properties of the mRNA-NPs: first, by concentrating numerous mRNAs on an iron-oxide nanoparticle with defined stoichiometry, the mRNA-NPs behave as hotspots of local translation activity that continuously produce nascent proteins. Second, the mRNA-based nanoparticles can be manipulated by magnetic forces to generate localized spatial patterns. An exciting perspective of this work concerns the implementation of mRNA-NPs in cells and organisms where the spatiotemporal control of RNA translation is key to determine cell fate behaviour.

Over the past years, several studies have identified how molecular players, such as cis-acting RNA localization elements, RNA-binding proteins, and the transport of RNA-granules by molecular motors along cytoskeletal fibers, that can act collectively to specifically regulate subcellular protein synthesis(*1*). However, the mechanisms regulating the dynamic behavior and spatial positioning of the synthetized proteins upon translation termination and protein release remain elusive. To remain localized in a subcellular area, specific mechanisms are used by cells to counterbalance the diffusion and spreading in space of the newly synthetized proteins. For instance, protein anchoring to a localized scaffold at the site of translation, and reduced mobility due to geometrical constraints or molecular crowding, may confine proteins at the site of translation. Interestingly in our study, by monitoring both local translation on the mRNA-NPs and nascent fluorescent protein synthesis, we found that mRNA can act as an anchoring molecule that can spatially retains N-terminal folded protein domains during translation. By examining the local translation of magnetic assemblies of mRNA-NPs coding for ABD, we found that the spatial retention of ABD on the mRNA-NP area was required to localize F-actin patterns (**Figures 3 and S6**). In addition, F-actin local assemblies can, in turn, modify the localization of the mRNA-NPs positioning; providing an interesting feed-forward mechanism between local translation-based mRNA localization mechanisms and cytoskeleton self-organization (**Figure 4**). In this picture, a temporary spatial retention of ABD renders the mRNA nanoparticle interactive with its own protein and functional partners (**Figure 5**). Experiments performed in bacteria and yeast underlie that that protein complexes can assembled while sub-unit are synthetized by the ribosome machinery co-translationally in cells(*43*–*45*). In additions, recent studies discovered novel functional roles of the non-coding parts of RNAs as scaffolds to promote protein-protein interactions, with the 3’ UTRs of mRNAs that can recruit proteins to the site of translation to determines subcellular protein localization(*46*), or with long non-coding RNA that can be a platform for post-translational functions(*47*). Here, we show that the coding sequence of mRNAs, in addition to providing genetic information, can also act as a scaffold molecule to localize protein-protein interactions between protein domains that fold during translation and binding protein partners (**Figure 5**).

**Figure 5:**
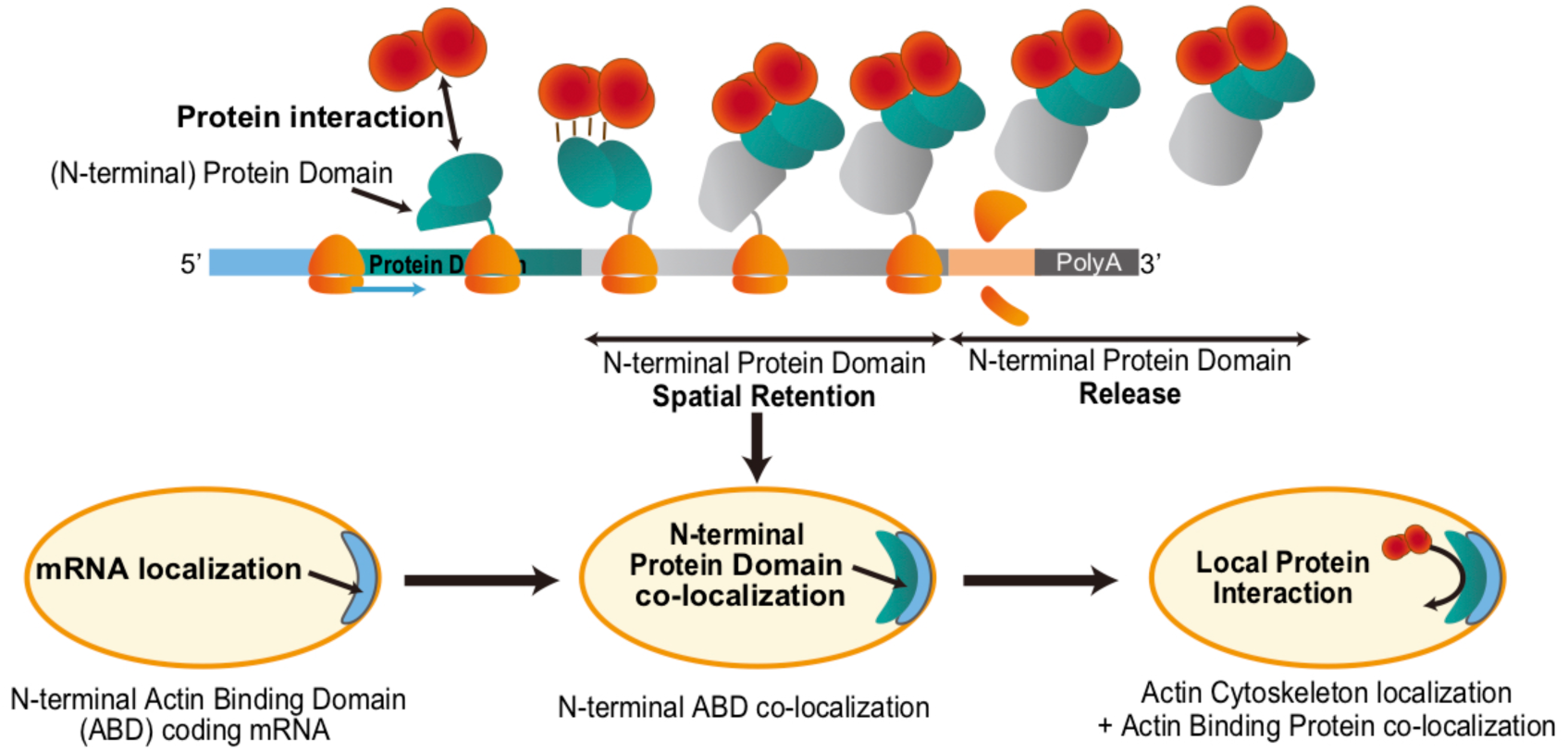
Proposed model of mRNA as translation-coupled scaffold with anchoring function to spatially constrain protein interactions. Spatially contrained protein interactions could be co-translationally mediated by the spatial retention of nascent protein domain on localized mRNA. For instance, the N-terminal positioning of actin binding domain sequence on localized mRNA leads to the spatial retention of the protein domain during translation and eventually promotes actin filament localization.

The mechanism of spatial retention of protein domains during local translation may have relevant physiological implications. Given the modular organization of protein domain structures, a temporary local concentration of functional modules that are produced earlier during translation, can be generated. Numerous Acting Binding Proteins have ABDs at their N terminals(*38*–*41*), suggesting the possibility of spatial and temporal retention of the ABDs on mRNA templates with a duration defined by the extent of the downstream mRNA translation process. For instance in humans, the Utrophin mRNAs and proteins are both known to be localized at neuromuscular junctions in mouse muscle cells and the long Utrophin mRNA sequence (9537 bases which 13-fold longer than GFP) may thus serve as scaffold to spatially retains ABD(*38, 39*).

Our *in vitro* local translation model allowed us to identify a possible anchoring function of mRNA occurring co-translationally and mediated by the spatial retention of N-terminal domain during translation (**Figure 5**). Further investigations need to be performed to know if this anchoring property could occur *in vivo* to enable localized protein-protein interactions by creating enriched concentration of nascent functional protein domains and therefore by participating in regulating local translation.

## Methods

### Plasmid construction and dsDNA template preparation

The genes for dsDNA templates were cloned into pT7CFE1-CHis cloning vector (pCFE, Thermo Fisher Scientific). Phusion high-fidelity DNA polymerase (New England Biolabs) was used for polymerase chain reaction (PCR). The sequences of oligo DNA primers and template gene for the construction were described in **Supplemental Table 1**.

GFP-ABD gene was amplified by PCR from pET28-GFP-hUtrophin(*48*), using the following oligonucleotides primers: *NdeI_eGFP fwd* and *hUtr_XhoI rev*. GFP-ABD was then cloned into pCFE between the NdeI and XhoI sites to construct the pCFE-GFP-ABD. mCherry gene was amplified by PCR from pET28-hUtrophin-mCherry(*48*) with *BamHI_mCherry fwd* and *mCherry_XhoI rev* primers. The gene was then replaced with ABD on pCFP-GFP-ABD between the BamHI and XhoI sites to construct the pCFP-GFP-mCherry. To construct pCFE-GFP-122aa a gene of 122 amino acids (a.a) from mWASP_VCA gene was amplified by PCR with *NcoI_122_a.a. fwd* and *122_a.a._XhoI rev* primers. The gene was then replaced with ABD on pCFP-GFP-ABD between the NcoI and XhoI sites. To construct pCFE-GFP-375aa-mCherry, GFP gene or 375aa gene was amplified from pCFE-GFP-ABD or synthetic gene of hUtrophin rods region (262-636) by PCR with *NdeI_eGFP fwd* and *eGFP_XbaI_hUtrn262 rev* or *hUtrn262 fwd* and *hUtrn636_BamHI rev* primers respectively. The two PCR products were then mixed and amplified by PCR with *NdeI_eGFP fwd* and *hUtrn636_BamHI rev* to fuse ABD and GFP genes. The PCR product was cloned into pCFE-GFP-mCherry between the NdeI and BamHI sites. To construct pCFE-ABD-GFP, ABD gene or GFP gene was amplified from pCFE-GFP-ABD by PCR with *NdeI_hUtrophin fwd* and *hUtr_link1 rev* or *link1_eGFP fwd* and *eGFP_XhoI rev* primers respectively. The two PCR products were then mixed and amplified by PCR with *NdeI_hUtrophin fwd* and *eGFP_XhoI rev* to fuse ABD and GFP genes. The PCR product was cloned into pCFE between the NdeI and XhoI sites. And all the sequences were verified.

DNA templates consist of T7 promoter, EMCV IRES sequence, gene of interest and 120 mer polyA sequence were amplified by PCR with *TAP_T7G3C fwd* and *3’UTR_120A rev* primers from the mixture of pCFE plasmids and *pCFE_3UTR15T* primer. PCR products were treated with DpnI (NEW ENGLAND BioLabs Inc) to degrade pCFE plasmids and purified with QIAquick PCR Purification Kit (QIAGEN). The quality and length of DNA templates were analyzed with agarose gel assay.

### In vitro transcription from DNA templates and Biotinylation of mRNA

The DNA templates were *in vitro* transcribed into mRNA with MEGAscript T7 Transcription Kit (Thermo Fisher Scientific) and purified with RNeasy Mini Kit (QIAGEN). For transcription of Cy5-labeled mRNA 7.5mM CTP was substituted with the mixture of 3.75 mM CTP and 0.25 mM 5-Propargylamino-CTP-CY5 (Jena Bioscience GmbH). The 3’ end of the mRNA were ligated with pCp-Biotin (Jena Bioscience GmbH) by T4 RNA Ligase (10 U/µL, Thermo Fisher Scientific) with following process. 1 µM of mRNA (or Cy5-labeled mRNA) and 50 µM of pCp-Biotin were mixed with 1 U/µL T4 RNA Ligase in ligation buffer and incubated for 2 hours at 16 °C and then purified with RNeasy Mini Kit. Both mRNA and incorporated Cy5 concentration were measured with NanoDrop 2000 spectrophotometer (Thermo Fisher Scientific).

### Conjugation of mRNA to nanoparticles

The 0.6-2.0 µg of biotinylated mRNAs (supplemented with Cy5-labeled biotinylated mRNAs) were incubated with 9.2×10^−16^ moles (for local translation) or 2.76×10^−14^ moles (for clustering) of 200 nm streptavidin coated magnetic nanoparticles (NP, Bio-Adembeads streptavidin 200 nm, Ademtech) for 45 minutes at 4 °C in high salt buffer [0.5 M NaCl, 20 mM Tris-HCl (pH 7.6)]. The mRNA conjugated NPs (mRNA-NPs) were washed with low salt buffer [0.15 M NaCl, 20 mM Tris-HCl (pH 7.5)]. The stoichiometries of conjugation were determined by the optimized amount of each mRNA for translation efficiency (**Supplementary Fig. 1a**). The conjugation efficiency was about 33% to 50% for the experiments using a stoichiometry of 800-1600:1 (mRNA:NP).

### Extract-in-Oil droplets formations and inclusion in cover slide chambers

5 µL of *HeLa* Cell Extract, 1 µL of Accessory protein and 2 µL of reaction mix (from 1-Step Human Coupled IVT Kit-DNA, Thermo Fisher Scientific) were mixed with 2 µL of mRNA-NPs and immediately encapsulated in droplets by water-in-oil emulsion process. The process has been previously described and allows the formation of active actin meshworks in droplets(*48*). PHS-PEO-PHS block copolymer (Arlacel P135) was first dissolved in mineral oil (0.4 mg/mL). The HeLa cell extract containing mRNA-NPs was then added to the block copolymer solution (3 % (v HCE/v Oil)) at room temperature (**Supplementary Fig. 1b**). The mixture is tapped five-to-six times to generate polydisperse extract-in-oil droplets. The mechanical dispersion of the biphasic solution formed micrometer-sized extract-in-oil droplets within few seconds. The emulsion was immediately enclosed in a handcrafted slide chamber and sealed with UV-sensitive photopolymer, Norland Optical Adhesive 60 (NOA60, Norland Products) followed by 30 seconds of 365 nm UV light exposure only on NOA60 (**Supplementary Fig. 1b**). The handcrafted slide chamber was produced with following processes. 2 × 20 × 0.15 mm sliced paper tape were attached on 24 × 32 mm cover glass (VWR) to make three edges of the chamber. Then the tree inner edges were caulked with NOA60 and covered by 22 x 22 mm cover glass (VWR) and followed by 5 minutes of UV light exposure to fix the chamber slide (**Supplementary Fig. 1b**).

## Imaging and Data Analysis

Fluorescence imaging of locally translated proteins and Phalloidin-568 (**Figure 2 and 3d**) was performed using an IX81 microscope (Olympus) and 60x oil immersion objective (PlanApo, NA 1.42), equipped with an EM-CCD camera (electron multiplying CCD, C9100-13, Hamamatsu, Corporation), and a LED system of illumination (Spectra X, Lumencor) in a microscope incubator (Oko-Lab) to keep the temperature of the slide chamber at 30 °C during time-lapse imaging. Microscope settings and time-lapse imaging functions were controlled using Micro-manager on ImageJ. Image analysis was performed using ImageJ and R. Confocal microscopy was performed (for **Figure 3 and 4**) with a Zeiss LSM 710 META laser scanning confocal microscope using 63x objective (PlanApo, NA 1.4) to imaging actin binding proteins translation and actin meshworks. Image analysis was performed using LSM Software Zen 2009 and ImageJ.

### Monitoring translation from asymmetrically localized mRNA-N

To monitoring local translation in droplets, 1.5×10^−12^ (or 0.4×10^−12^) moles of biotinylated mRNA(GFP) [or mRNA(GFP-mCherry)] were mixed with streptavidin coated nanoparticles to produce mRNA-NP. The droplets including 0.46 nM of mRNA-NP and *HeLa* cell extract were imaged with time-lapse microscope in the slide chamber for 2∼4 hours at 30 °C besides a permanent NdFeB magnet (Q-15-04-04-MN, Supermagnete) in a distance ranging from 5 to 10 mm from the droplets. The mean intensities of whole droplet and selected area were normalized by subtracting background intensities. The mean intensities of whole droplets area were divided by droplet diameters to calculate the relative concentrations of droplets. For the bleach correction of GFP and mCherry mean intensities, the fluorescent values were divided by the bleaching rate that was computed by fitting bleach curves obtained from dedicated experiments of translated GFP-X proteins performed with the same observation condition. The 4-parameter logistic curve fitting to the time-series of the fluorescent intensities were performed with Nonlinear Least Squares function of R with formula F(logT)=Base+(Max-Base)/ (1+10^(n*(log Imax(counter edge)/2-log T).

### F-actin meshwork morphology modulated by local translation of mRNA-NP

1.7×10^−12^ moles of biotinylated mRNA(GFP-ABD) or mRNA(ABD-GFP) and corresponding biotinylated Cy5-labelled mRNA was used for mRNA-NP conjugation. The HeLa cell extract droplets including 0.46 nM of mRNA-NPs, 6 µM of G-Actin from rabbit skeletal muscle (AKL95, Cytoskeleton Inc.) and 100 nM of Alexa Fluor 568 Phalloidin (only for samples with phalloidin labelled images, Life Technologies) in the slide chamber were incubated and imaged besides a NdFeB magnet (or without magnet) for 2-3 hours with confocal microscope.

## Supporting information

## Authors’ contributions

S.K, D.O.W, H.S, and Z.G conceived and designed the experiments. S.K performed the experiments. S.K and Z.G analyzed the experiments. D.O.W, H.S contributed materials/analysis tools. S.K and Z.G wrote the manuscript and all authors commented on it.

### Acknowledgements

We acknowledge the members of the Biophysical Chemistry group of ENS for fruitful discussions. We thank Dr. Kei Endo for reading carefully the manuscript. The authors also thank I. Oomoto, C. Parr, A. Hubstenberger, and Y. Fujita. S.K was supported by a fellowship “bourse du gouvernement français”. This work was supported by the HFSP Program Grant (RGP0050/2014) to D.O.W, H.S, and Z.G; by the CNRS to Z.G; and by the Ecole Normale Supérieure to Z.G.

