## Supplementary material for "Nanoparticle-based local translation reveals mRNA as translation-coupled scaffold with anchoring function"

### Index of SUPPLEMENTARY INFORMATION:

**8 Figures, 1 Tables, 7 Supplementary movies**

**Supplementary Discussion and Experimental Procedures:**

1. **Supplementary Figures**
2. **Supplementary Table**
3. **Legends for Supplementary Movies**

### 1. Supplementary Figures

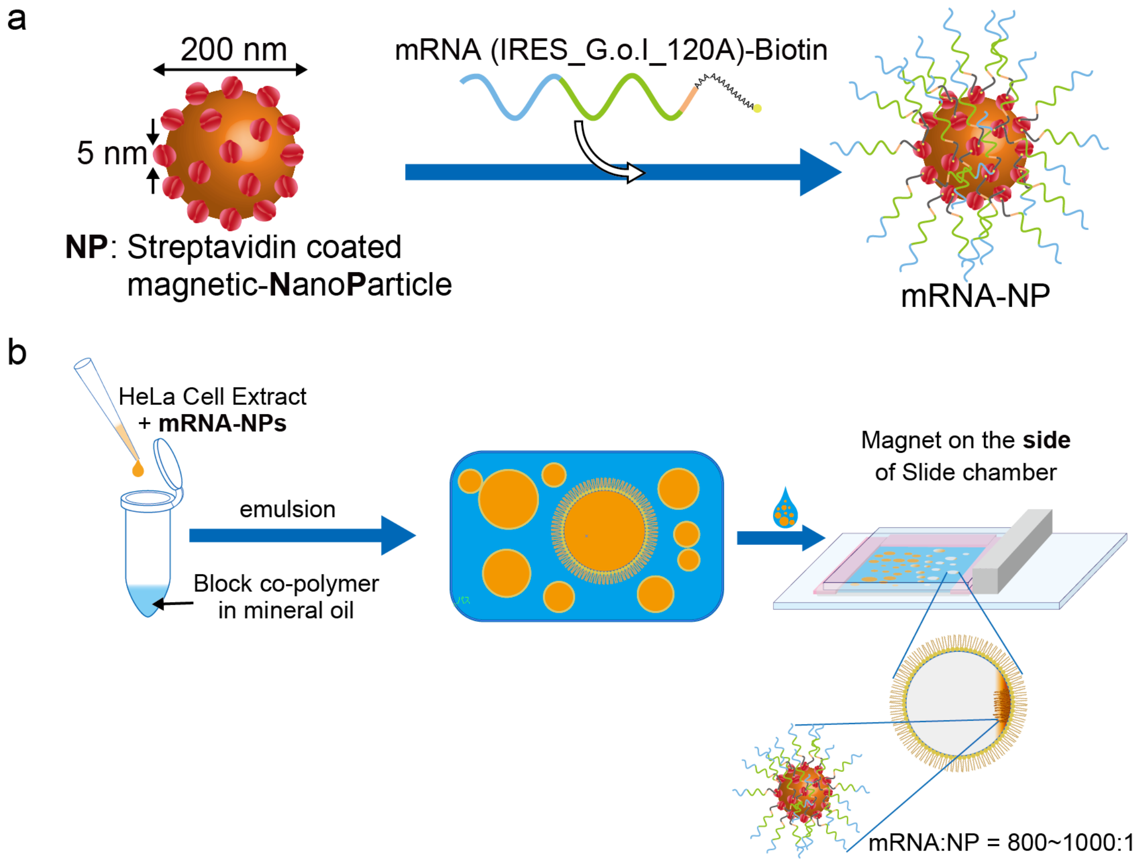


**Supplementary Figure 1. Translation of mRNA-nanoparticle conjugates in *HeLa* cell extracts.**

Experimental scheme of mRNA-NP local translation and clustering. **a**, Streptavidin coated magnetic-nanoparticles (NPs, 200 nm) are iron oxide nanoparticles covered with 5 nm streptavidin tetramers. Schematic of the mRNA sequence (IRES_G.o.I_120A)-biotin consisting of an Internal Ribosome Entry Site, a Gene of Interest, and 120 mer adenines and biotinylated cytosine at the 3-prime end. The 3' end of biotinylated mRNA links with NP by the tight streptavidin-biotin interaction ($K_d \sim 10^{-14}$). **b**, The mRNA-NPs dispersed in *HeLa* cell extract are added to a mixture of mineral oil with block co-polymers in order to be confined into micrometer scale droplet. The *HeLa* extract droplets are then enclosed into an observation chamber. The conjugation ratio of mRNA to NP is optimized for both mRNA-NP patterns and translation efficiency. For generating asymmetric mRNA-NP localization, a permanent magnet is placed besides the slide chamber.

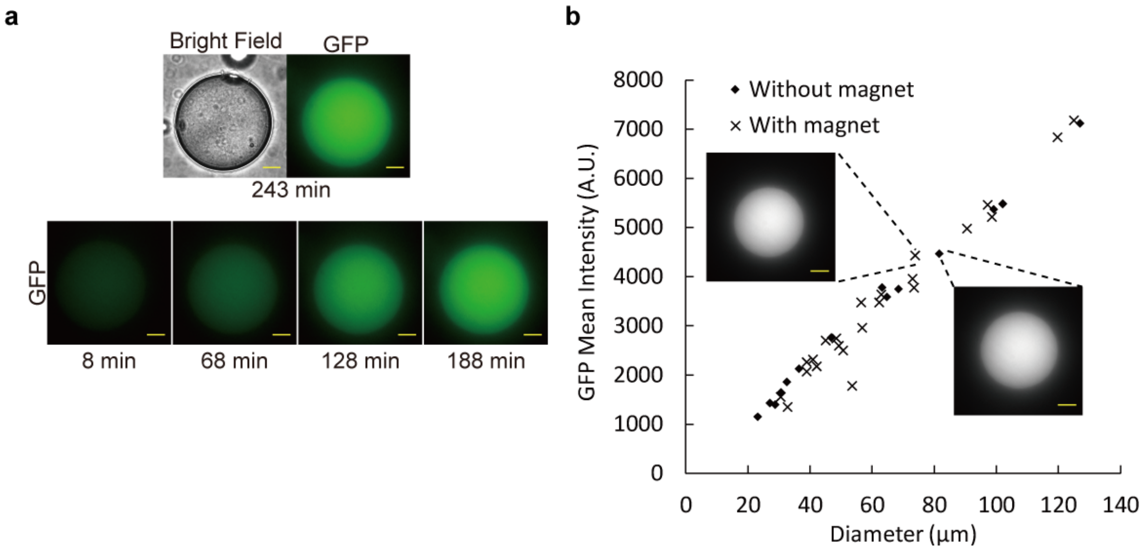


**Supplementary Figure 2. Translational efficiency of the mRNA-NPs**

**a**. The representative images of continuous GFP production after the translation of conjugated (mRNA(GFP)-NPs). The translation reaction was performed in confined *HeLa* extracts without magnet. **b**, Plots of the GFP mean intensities (after 30 °C, 4 hours incubation) as function of the droplet diameter in the presence and the absence of magnetic field. Scale bar, 20 µm. Each data point is corresponding to the data from Figure 2c.

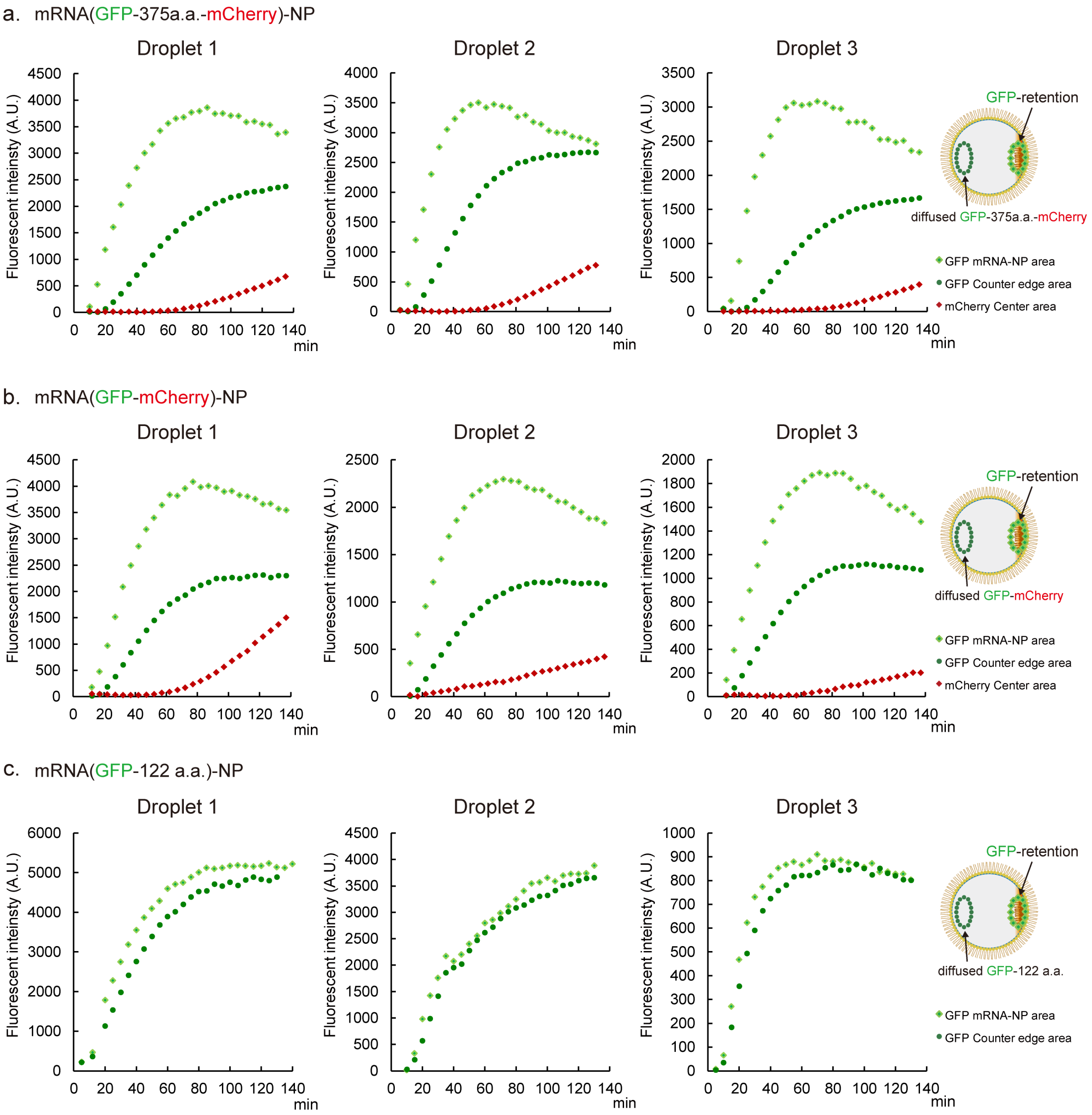


**Supplementary Figure 3. Time courses of translation of asymmetrically localized mRNA-NPs**

Time dependent evolution of fluorescent intensity of GFP and mCherry in selected regions of the droplets (one frame every 5 minutes) for three respective droplet experiments. The nanoparticles were conjugated to mRNA(GFP-375a.a.-mCherry) (**a**); mRNA(GFP-mCherry) (**b**); or mRNA(GFP-122a.a.) (**c**).

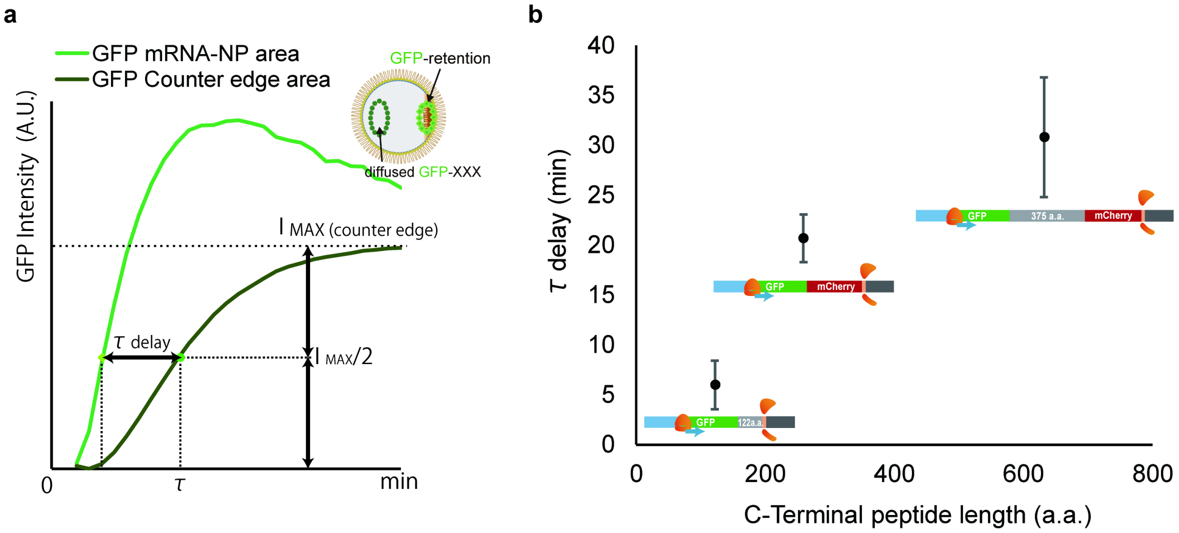


c

| N-terminal | C-terminal | bases | a.a. | Tdelay (min) | SD |
|---|---|---|---|---|---|
| GFP- | **122 a.a.** | **369** | **122** | **6** | **2.45** |
| | **mCherry** | **780** | **259** | **20.7** | **2.39** |
| | **375 a.a.-mCherry** | **1899** | **632** | **30.8** | **5.98** |

**Supplementary Figure 4. Estimation of the time difference in fluorescence appearance between the localized GFP signal on RNA-NPs and the GFP diffuse fraction in the droplet**

**a**. Time delay τ calculation and definition. The time series of GFP intensities in the both area of droplets were fitted to 4-parameter logistic curves. The time delay τ is defined as the time difference between the mean intensity measured in the mRNA-NP area and in the counter edge area (taken at the half of intensity max in counter edge area ($I_{MAX(counter\ edge)}/2$)). **b**,**c.** Plot of the time delay τ as a function of the downstream C-terminal peptide length (a.a.; amino acids, SD; standard deviation of three different experiments).

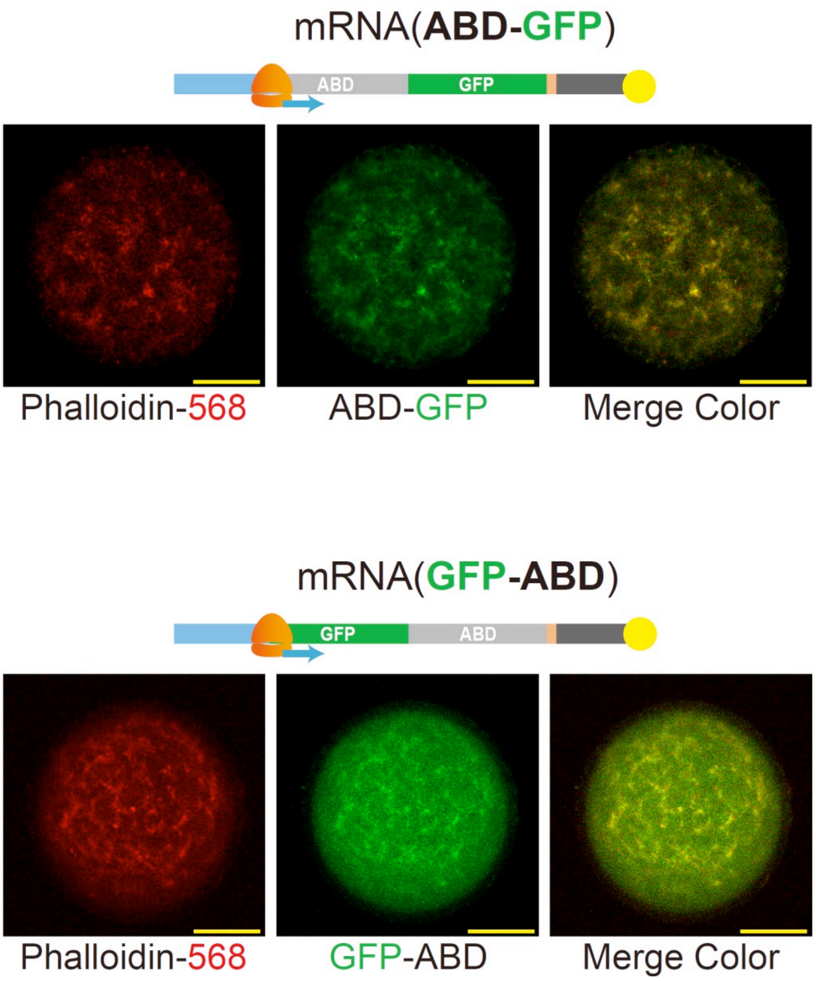


**Supplementary Figure 5. Actin filament meshwork induced by the translation of mRNA(ABD-GFP) or mRNA(GFP-ABD)**

Representative confocal fluorescence microscopy images of actin filament meshwork and ABD localization observed with phalloidin-568 (F-actin marker), and ABD-GFP or GFP-ABD. mRNA(ABD-GFP)-biotin or mRNA(GFP-ABD)-biotin without NPs were translated with 6 μM G-actin and 100 nM phalloidin-568 in *HeLa* cell extract droplets (30 °C for 5 hours). Scale bar, 20 μm.

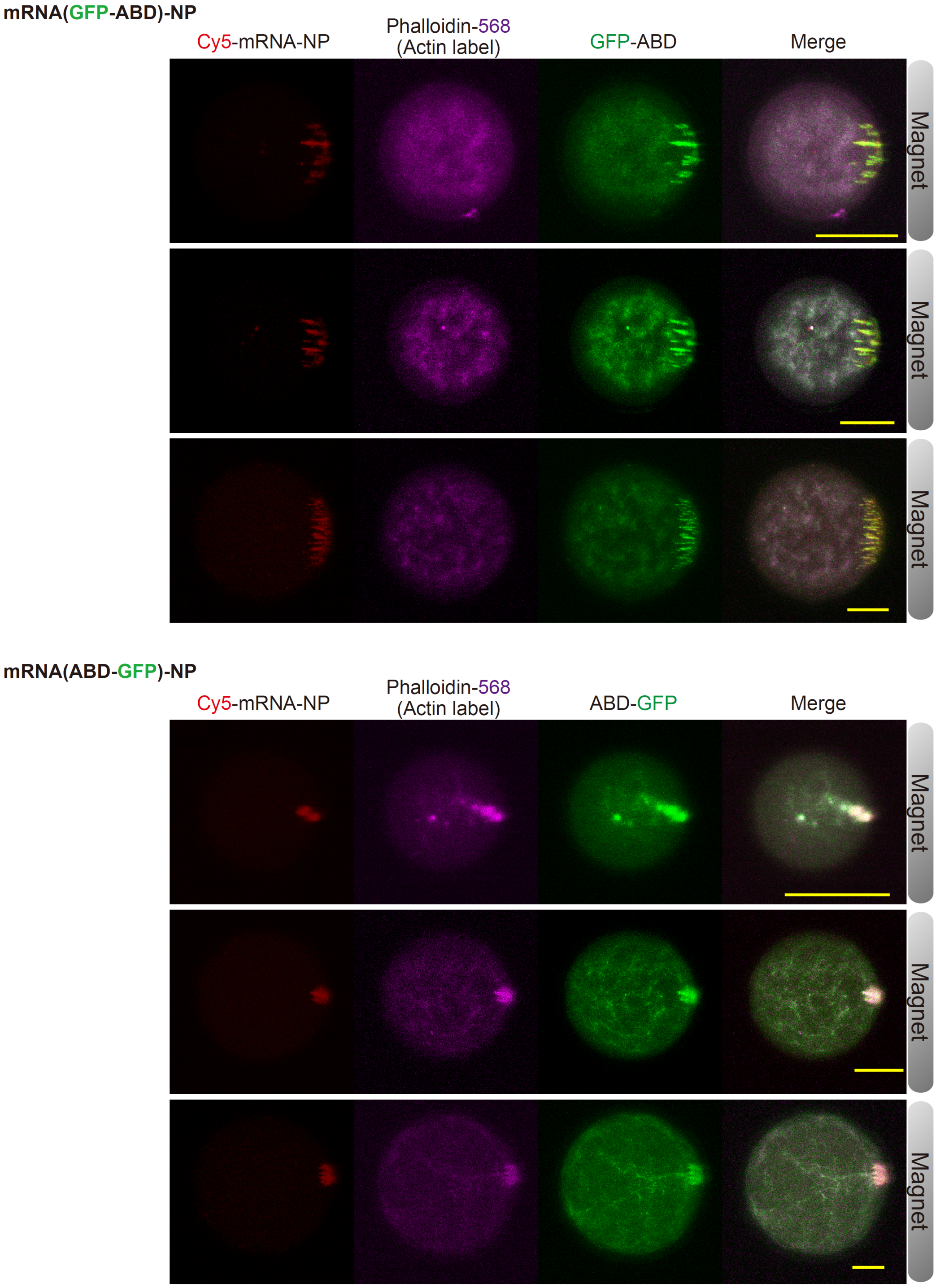


**Supplementary Figure 6. Spatial control of F-actin meshwork by localizing mRNA-NP translation**

Examples of confocal fluorescence microscopy images of Cy5-mRNA- NPs, phalloidin-568 (F-actin marker), and GFP-ABD (**top**) or ABD-GFP (**bottom**), after 3 hours of translation reaction. Scale bar, 20 μm.

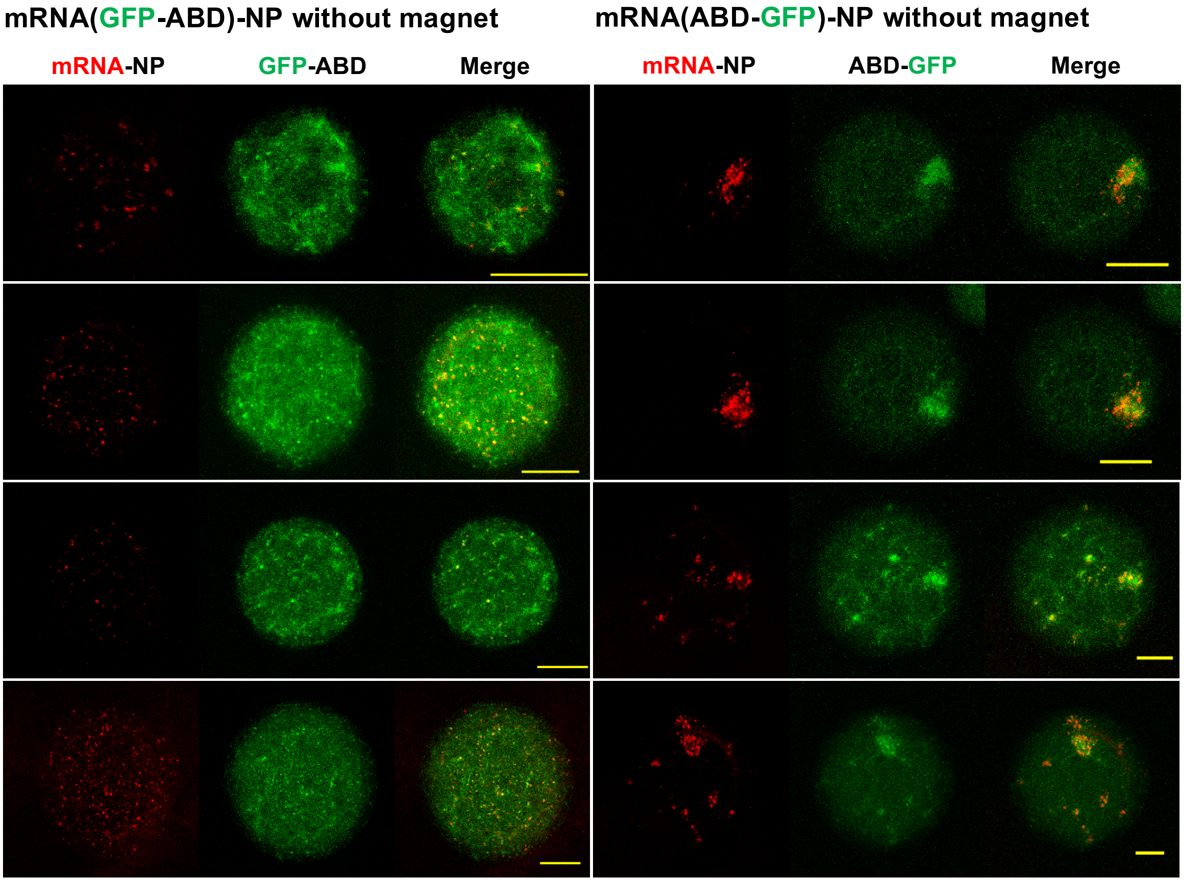


**Supplementary Figure 7. The pattern of F-actin meshwork depends on the position of ABD domain on mRNA (N-terminal or C-terminal position) during mRNA-NP translation**

Examples of confocal fluorescence microscopy images of Cy5-mRNA-NPs and GFP-ABD (**left**) or ABD-GFP (**right**), after 3 hours of translation reaction. Scale bar, 20 μm.

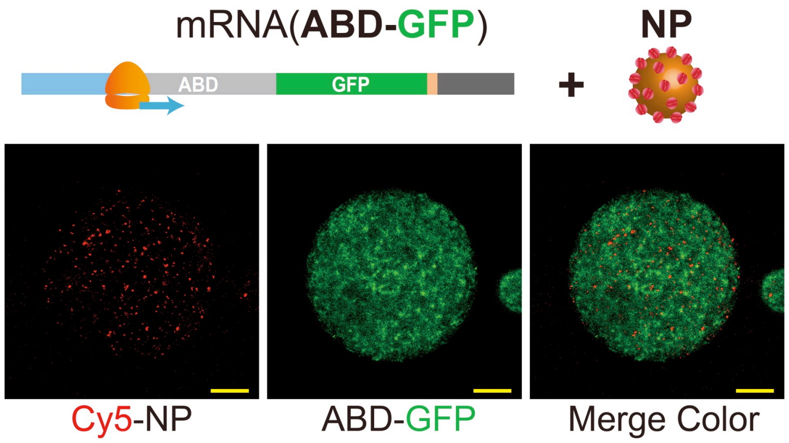


**Supplementary Figure 8. Translation of non-biotinylated mRNA(ABD-GFP) in the presence of unconjugated NPs**

Examples for confocal fluorescence microscopy images of Cy5-NPs, phalloidin-568 (F-actin marker), and GFP-ABD (after 3 hours incubation at 30 °C). mRNA(ABD-GFP) without biotin were mixed with Cy5-labeled NPs in *HeLa* cell extract. Scale bar, 20 μm.

### 3. Supplementary Table

The oligo DNA sequences used for the experiments

| Primer name | Sequence |
|---|---|
| NdeI_eGFP fwd | 5’-gggcacCATATGgtgagcaagggcgagga-3’ |
| eGFP_XhoI rev | 5’-ggtgcaCTCGAGcttgtacagctcgtccatgcc-3’ |
| BamHI_mCherry fwd | 5’-cagctcGGATCCgtgagcaagggcgaggaggataac-3’ |
| mCherry_XhoI rev | 5’-gttCTCGAGcttgtacagctcgtccatgccg-3’ |
| NdeI_eGFP fwd | 5'-GGGCACCATatggtgagcaagggcgagga |
| eGFP_XbaI_hUtrn262 rev | 5'-GAGTGTCTCTACCTCACGGATGGCgctgcctccactaccgccTCTAGACTTGTACAGCTCGTCCATGCC |
| hUtrn262 fwd | 5'-GCCATCCGTGAGGTAGAGACACTC |
| hUtrn636_BamHI rev | 5'-cggcgtggatcctccactacctccAGTCACCTGGTTGGAGGAATCTTC |
| NcoI_122_a.a. fwd | 5'-ggactgCCATGGaccatcaagttccagc |
| 122_a.a._XhoI rev | 5'-GGCGAActcgagGTCTTCCCACTCATCATCATCC |
| NdeI_hUtrophin fwd | 5’-gctcagCATATGgccaagtatggagaacatgaag-3’ |
| hUtr_link1 rev | 5’-GGATCCTCCACTACCTCCGCTACCgtctatggtgacttgctgaggtagc-3’ |
| link1_eGFP fwd | 5’-GGTAGCGGAGGTAGTGGAGGATCCgtgagcaagggcgaggag-3’ |
| hUtr_XhoI rev | 5’-gccCTCGAGgtctatggtgacttgctgaggtagc-3’ |
| TAP_T7G3C fwd | 5’-cagtgaattgtaatacgactcactatagggc-3’ |
| 3'UTR_120A rev | 5’-ttttttttttttttttttttttttttttttttttttttttttttttttttttttttttttttttttttttttttttttttttttttttttttttttttttttttttttttttttttttttcctactcaggctttattca-3’ |
| pCFE_3UTR15T | 5’-tttttttttttttttcctactcaggctttattcacagtcagatctcagtggtgg-3’ |
| 375 a.a. linker from hUtrophin rods region (262-636) | 5’-GCCATCCGTGAGGTAGAGACACTCccaaggaaatataaaaaagaatgtgaagaagaggcaattaatatacagagtacagcgcctgaggaggagcatgagagtccccgagctgaaactcccagcactgtcactgaggttgacatggatctggacagctatcagattgcgttggaggaagtgctgacctggttgctttctgctgaggacactttccaggagcaggatgatatttctgatgatgttgaagaagtcaaagaccagtttgcaacccatgaagcttttatgatggaactgactgcacaccagagcagtgtgggcagcgtcctgcaggcaggcaaccaactgataacacaaggaactctgtcagacgaagaagaatttgagattcaggaacagatgaccctgctgaatgctagatgggaggctcttagggtggagagtatggacagacagtcccggctgcacgatgtgctgatggaactgcagaagaagcaactgcagcagctctccgcctggttaacactcacagaggagcgcattcagaagatggaaacttgccccctggatgatgatgtaaaatctctacaaaagctgctagaagaacataaaagtttgcaaagtgatcttgaggctgaacaggtgaaagtaaattcactaactcacatggtggtcattgttgatgaaaacagtggtgagagtgctacagctatcctagaagaccagttacagaaacttggtgagcgctggacagcagtatgccgttggaCTGAAGAACGCTGGAATAGGTTACAAgaaatcaatatattgtggcaggaattattggaagaacagtgcttgttgaaagcttggttaaccgaaaaagaagaggctttaaataaagtccagacaagcaacttcaaagaccaaaaggaactaagtgtcagtgttcgacgtctggctattttgaaggaagacatggaaatgaagcgtcaaacattggatcagctgagtgagattggccaggatgtgggacaattacttgataattccaaggcatctaagaagatcaacagtgactcagaggaactgactcaaagatgggattctttggttcagagactaGAAGATTCCTCCAACCAGGTGACT-3’ |
| 122 a.a. linker from VCA_mWASP gene | 5'-atggaccatcaagttccagctccttcaggaaacaaagcagctcttttggatcaaattagagagggtgctcagctgaaaaaagtggaacagaatagccggcccgtgtcctgctcaggaagggatgcgcttttagaccagatacgacagggcattcaattgaaatctgtgtctgatggccaagagtccacaccaccaacacccgcacccacttcaggaattgtgggtgcgctgatggaagtgatgcagaanaggagcanagccattcattcctcagatgaagatgaagacgatgatgatgaagaagattttgaggatgatgatgagtgggaagac-3' |

### 4. Legends for Supplementary movies

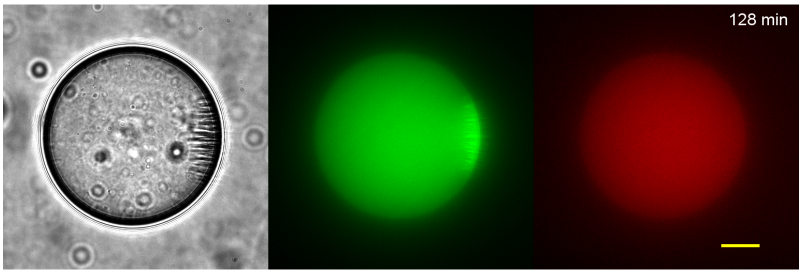


**Supplementary Movie 1.**

**Translation from asymmetrically localized mRNA(GFP-mCherry)-NPs in droplet.**

Left panel: Bright field observation of mRNA-NP string-like assemblies. Middle, and right panels: Time dependent production of GFP (middle panel) and mCherry (right panel). The direction of the magnetic forces was directed towards the right. Epifluorescence microscopy, Scale bar: 20 µm, one frame every 15 minutes.

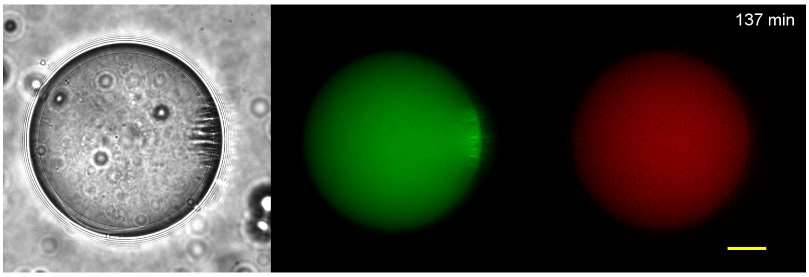


**Supplementary Movie 2.**

**Translation from asymmetrically localized mRNA(GFP-mCherry)-NPs in droplet.**

Left panel: Bright field observation of mRNA-NP string-like assemblies. Middle, and right panels: Time dependent production of GFP (middle panel) and mCherry (right panel). The direction of the magnetic forces was directed towards the right. Epifluorescence microscopy, Scale bar: 20 µm, one frame every 5 minutes.

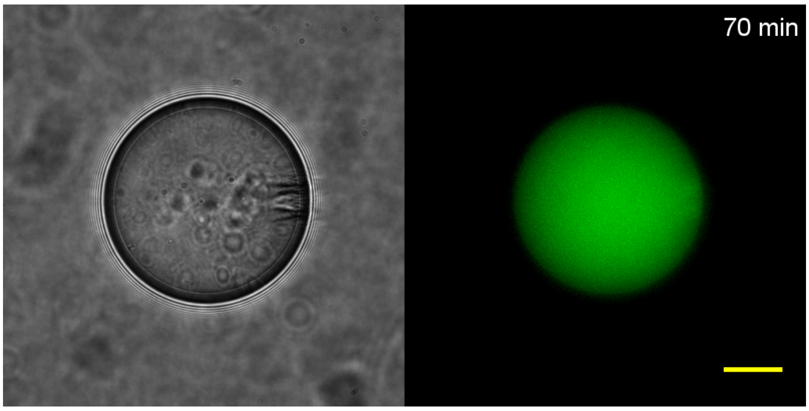


**Supplementary Movie 3.**

**Translation from asymmetrically localized mRNA(GFP-122aa)-NPs in droplet.**

Left panel: Bright field observation of mRNA-NP string-like assemblies. Right panels: Time dependent production of GFP. The direction of the magnetic forces was directed towards the right. Epifluorescence microscopy, Scale bar: 20 µm, one frame every 5 minutes.

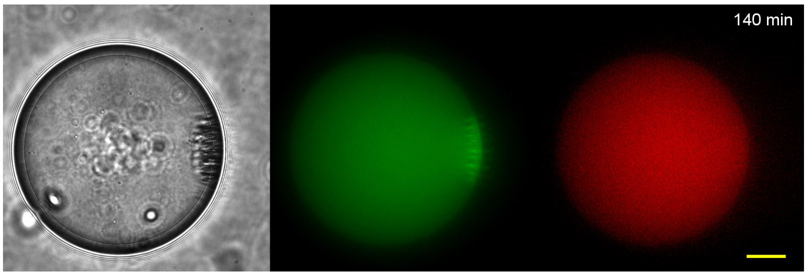


**Supplementary Movie 4.**

**Translation from asymmetrically localized mRNA(GFP-375aa-mCherry)-NPs in droplet.**

Left panel: Bright field observation of mRNA-NP string-like assemblies. Middle, and right panels: Time dependent production of GFP (middle panel) and mCherry (right panel). The direction of the magnetic forces was directed towards the right. Epifluorescence microscopy, Scale bar: 20 µm, one frame every 5 minutes.

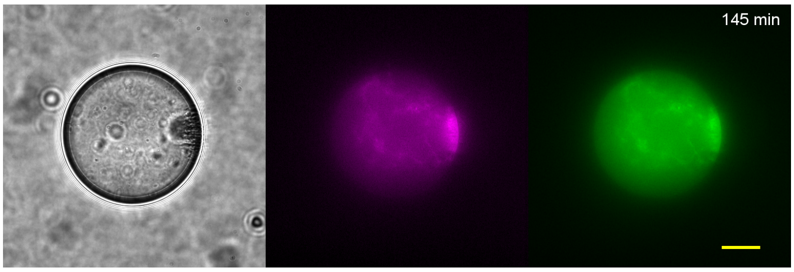


**Supplementary Movie 5.**

**Translation from asymmetrically localized mRNA(ABD-GFP)-NPs in droplet**

Left panel: Bright field observation of mRNA-NP string-like assemblies. Middle panel: Time dependent accumulation of Phalloidin-568 as label for F-actin. Right panel: Time dependent production of GFP. The direction of the magnetic forces was directed towards the right. Epifluorescence, Scale bar: 20 µm, one frame every 5 minutes.

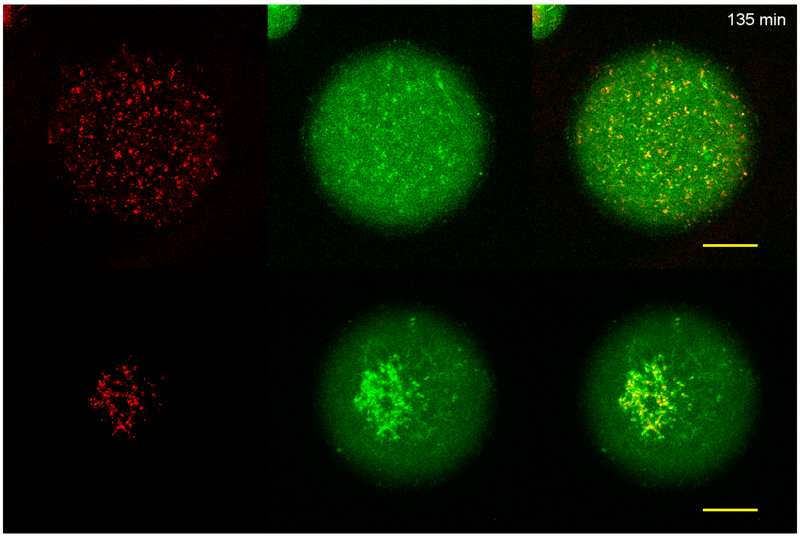


**Supplementary Movie 6.**

**The spatial retention of ABD on mRNA-NPs impacts the spatiotemporal dynamics of F-actin assemblies**

Translation of Cy5-labelled mRNA(GFP-ABD)-NPs (**top three panels**) and Cy5-labelled mRNA(ABD-GFP)-NPs performed in absence of magnetic forces (**bottom three panels).** Left panels: Cy5 observation of mRNA-NPs. Middle panels: Time dependent production and localization of GFP. Right panels: Merge color images. Confocal fluorescence microscopy, Scale bar: 20 µm, one frame every 15 minutes.

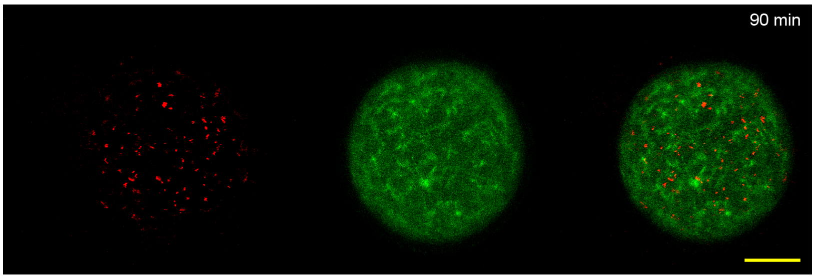


**Supplementary Movie 7.**

**Translation of non-biotinylated mRNA(ABD-GFP) in the presence of unconjugated Cy5-labelled NPs mRNA(ABD-GFP)**

Left panels: Cy5 observation of nanoparticles. Middle panels: Time dependent production and localization of ABD-GFP. Right panels: Merge color images. Confocal fluorescence microscopy, Scale bar: 20 µm, one frame every 15 minutes.
